# Predisposing factors of teenage pregnancy in the Uganda Lake Victoria Island and Mountain districts

**DOI:** 10.1101/482927

**Authors:** Richard Tuyiragize, Abel Nzabona, John Bosco Asiimwe, Christian Kakuba, John Mushomi, Fred Maniragaba

## Abstract

**Introduction:** There is a high teenage pregnancy in Lake Victoria Island and Mountain districts of Uganda. Teenage pregnancy is highly associated with abortions, infant and maternal mortality, high rate of unemployment, school failure and drop-outs and limited future career opportunities. This paper identifies and explains the factors influencing teenage pregnancy in Uganda Lake Victoria Island shoreline area and mountain districts.

**Methods:** The analysis focused on 405 girls aged 15–19 years, generated from the 2016 Uganda demographic and Health Survey. Odds ratios with 95% confidence interval and p-values were computed using appropriate logistic regression models to determine the presence and strength of associations between the teenage pregnancy and independent variables.

**Results:** Age, residence, secondary or higher education level, female-headed households, marital status (married), occupation, wealth index(rich quintile), and knowledge of ovulation cycle were found to have statistically significant associations with teenage pregnancy.

**Conclusion:** Increased age, rural residence, occupation(not working), and knowledge of ovulation cycle were statistically significant predisposing factors of teenage pregnancy in Uganda Lake Victoria Island shoreline area and mountain districts. Teenagers in these study areas should be provided with sexual education as well as teenage-friendly health services at health facilities that include a wide range of options, as well as medically accurate counselling and information could mitigate teenage pregnancy.

## Introduction

Teenage pregnancy and childbirth to women less than 20 years old continues to be a major global public health concern, affecting more than 16 million girls and young women worldwide(WHO, 2014). Teenage pregnancy is highly associated with abortions, infant and maternal mortality, high rate of unemployment, school failure and drop-outs and limited future career opportunities. As a result of increased awareness of the socioeconomic consequences of teenage pregnancy, researchers and policy makers have concluded that teenage pregnancy and childbearing is a serious problem (Gebregzabher, Hailu, & Assefa, 2018; Lillian & Mumbango, 2015; Gyan, 2013; Yakubu & Salisu, 2018; Ayanaw Habitu, Yalew, & Azale Bisetegn, 2018; Omoro et al., 2017).

According to the findings of the Uganda Demographic and Health Survey (UDHS) (UBOS, 2016), one out four (25%) girls aged 15 – 19 years have either a child or are pregnant, representing a 1% increase in teenage pregnancy rates over the previous 2011 survey,(UBOS, 2011). The highest prevalence of teenage pregnancy is in Lake Victoria Island districts at 48%,(UBOS, 2016). This shows that teenage pregnancy remains a major issue in the Lake Victoria Island districts. The low use of contraception has been associated with high fertility, which remains a public health concern that should be averted.

According to UDHS (2011) approximately 35% of girls drop out of school because of early marriage and 23% do so because of early pregnancy. Early childbearing carries particular risks, including dropping out of school, abandoning babies and obtaining illegal abortion that may result into death. Lillian & Mumbango (2015) revealed that teenage pregnancy and childbearing is a serious social problem that is linked to the spread of HIV/ AIDS, sexual abuse, neglect, and abortions as well as infant and maternal mortality. Their results showed that teenage pregnancy was influenced by generation, region, highest educational level, socio-economic status and cultural factors. Kabagenyi, Habaasa & Rutaremwa (2016) conducted a study to examine what influences teenager’s use of contraception among teenagers in Uganda. Their findings show that the key predictors were age at first birth, history of previous birth, current age, place of residence, education and socioeconomic status. The conclusion was great need to address barriers to use of contraception among young people. Use of contraception and improving access to the services is highly recommended to avert some of the unplanned births among these females(Babirye, Akulume, Kisakye, & Kiwanuka, 2018; Murphy-Erby, Stauss, & Estupinian, 2013).

According to UDHS(2016), the Eastern and East Central regions showed the highest rates of teenage pregnancy in Uganda with 30.1% and 31.6% respectively which is higher than the national figure. This is as a result of unsafe sexual practices. In addition to the unwanted/ early/ teenage pregnancies, these young people are also at a high risk of HIV infection and infection from other STIs (Lwihula, Outwater, & Nyamuryekung’e, 2006). There is need for an integrated approach to curb teenage pregnancy. Atuyambe et al (2008) concluded that pregnant teenagers in Wakiso district (Including Lake Victoria shoreline areas) lack basic needs like shelter, food and security. They also face relational problems with families, partners and the community. Several social factors such as religious beliefs, idleness and economic factors have been identified as factors contributing to early pregnancy and marriage (Gideon, 2013; Amoran, 2012; Donoghue, 1992; UNICEF, 2015; Jewkes, Vundule, Maforah, & Jordaan, 2001; Kaye, 2008; Mollborn, 2010).

There is, therefore, a need to sensitize the community and school personnel about adolescent reproductive health issues. In addition, adolescent friendly services need to be established or strengthened. Continuous in-service training for health workers with emphasis on counseling skills for young people is urgently needed. This paper identifies and explains the factors influencing teenage pregnancy in Uganda Lake Victoria Island shoreline area and mountain districts.

## Materials and Methods

The paper uses secondary data extracted from the 2016 Uganda Demographic and Health Survey (UDHS) dataset. The 2016 UDHS special areas include the Islands and shoreline districts(Kalangala, Mayuge, Buvuma, Namayingo, Rakai, Mukono and Wakiso) and mountains districts (Bundibugyo, Kasese, Ntoroko, Bukwo, Bulambuli, Kapchorwa, Kween, Kisoro, Sironko, Mbale, and Kaabong).

The UDHS used a multistage cluster sampling, whereby at first stage, a random sample of enumeration areas (EA), which are primary sampling units, was chosen from the census sampling frame. From the selected EAs, households were systematically drawn. Only women of reproductive age (15–49 years), in the selected households, were interviewed using a face-to-face questionnaire.

The questionnaire included variables on individual bio demographic factors, household characteristics, and sexual history. In our study, the main variable of interest was age at first birth or pregnancy of a woman. If yes, it was coded 1 and 0 otherwise, for women in the reproductive age-group. The explanatory variables included current age, education level, residence, household head (male or female), relationship to household head, marital status, religion, occupation/employment, social economic status (wealth index), knowledge of any contraception method, and knowledge of fertile period.

Data analysis was conducted at three stages to explore the predisposing factors of teenage pregnancy in Uganda Lake Victoria Island shoreline area and mountain districts. There was generation of descriptive statistics of demographic and socio-economic variables. Some independent variables were cross tabulated with teenage pregnancy to establish any potential associations. At the multivariate stage binary logistic regression was used, and both unadjusted and adjusted logistic regression findings presented. All the data were weighted to account for clustering and design effect. STATA 15 was used for the analyses.

## Results

### Characteristics of respondents

Majority of the teenagers were below 18 years (66.0%) and about 14.7% had their first birth at less than 15 years (Table 1). The percentage of teenagers with primary or no education (69.2%) more than doubled those with secondary or higher education. This implies that educating teenagers may have a significant effect in mitigating against teenage pregnancy. Majority of the teenagers (77.1%) were living in the rural areas, with their parents (47.5%), while one in three (33.4%) teenagers came from female-headed households, and were never married (77.2%).

**Table 1:** Characteristics of respondents

| Background characteristics | Number | Percentage |
| --- | --- | --- |
| **Age in years** |  |  |
| 15 | 86 | 21.0 |
| 16 | 92 | 22.8 |
| 17 | 90 | 22.2 |
| 18 | 82 | 20.4 |
| 19 | 55 | 13.6 |
| **Age at first birth** |  |  |
| Less than 15 years | 13 | 14.7 |
| 15 – 19 years | 75 | 85.3 |
| **Special areas** |  |  |
| Lake Victoria Island shoreline area districts | 38 | 9.3 |
| Mountain districts | 367 | 90.7 |
| **Residence** |  |  |
| Urban | 93 | 22.9 |
| Rural | 312 | 77.1 |
| **Education level** |  |  |
| None | 16 | 4.0 |
| Primary | 264 | 65.2 |
| Secondary + | 124 | 30.8 |
| **Sex of household head** |  |  |
| Male | 269 | 66.6 |
| Female | 135 | 33.4 |
| **Marital status** |  |  |
| Never married | 312 | 77.2 |
| Married | 81 | 20.0 |
| Ever married | 12 | 2.8 |
| **Religion** |  |  |
| Anglican | 136 | 33.5 |
| Catholic | 141 | 34.9 |
| Moslem | 56 | 13.8 |
| Other | 72 | 17.8 |
| **Occupation** |  |  |
| Working | 196 | 48.5 |
| Not working | 208 | 51.5 |
| **Wealth index** |  |  |
| Poorest | 42 | 10.5 |
| Poorer | 90 | 22.1 |
| Middle | 108 | 26.7 |
| Richer | 83 | 20.4 |
| Richest | 82 | 20.2 |
| **Knowledge of any FP method** |  |  |
| Knows FP methods | 392 | 97.0 |
| Knows nothing | 12 | 3.0 |

**Table 1: Continued**
| **Knowledge of fertile period (ovulation cycle)** |  |  |
| --- | --- | --- |
| During periods | 20 | 4.9 |
| After periods | 160 | 39.7 |
| Middle of periods | 42 | 10.3 |
| Before periods | 43 | 10.6 |
| Any time | 31 | 7.7 |
| Don't know | 109 | 26.8 |

There were only 20.2% of the teenagers in the richest wealth quintile. Asked about their religious affiliation, 34.9% of the teenagers were Catholics, while 33.5% were Anglicans and the majorities were not using contraception (87.9%). Whereas 97.0% of the teenagers had knowledge of family planning methods, 60.3% did not know the fertile period (ovulation cycle).

### Prevalence of teenage pregnancy

Teenage pregnancy in Lake Victoria Island districts was twice (48.7%) that of Mountain districts (24.3%), as well as twice the national figure of 25%. Table 2 indicates that teenage pregnancy was higher for teenagers in rural areas than urban ones, and the association was statistically significant (p=0.040).

**Table 2:** Prevalence of teenage pregnancy by background characteristics

| Background characteristics | No pregnancy | Pregnancy | Total |
| --- | --- | --- | --- |
| **Residence** |  |  |  |
| Urban | 82.3 | 17.7 | 93 |
| Rural | 70.9 | 29.1 | 312 |
| | $\chi^2 = 6.43$ | $p=0.040$ | |
| **Special areas** |  |  |  |
| Lake Victoria Island shoreline area districts | 51.7 | 48.3 | 38 |
| Mountain districts | 75.7 | 24.3 | 367 |
| | $\chi^2 = 13.6$ | $p=0.000$ | |
| **Education level** |  |  |  |
| None | 66.2 | 33.8 | 16 |
| Primary | 69.6 | 30.4 | 264 |
| Secondary + | 82.2 | 17.8 | 125 |
| | $\chi^2 = 10.4$ | $p=0.039$ | |
| **Sex of household head** |  |  |  |
| Male | 68.3 | 31.7 | 270 |
| Female | 83.8 | 16.2 | 135 |
| | $\chi^2 = 14.8$ | $p=0.000$ | |
| **Marital status** |  |  |  |
| Never married | 91.1 | 8.9 | 312 |
| Married | 8.8 | 91.2 | 81 |
| Ever married | 15.9 | 84.1 | 12 |
| | $\chi^2 = 294$ | $p=0.000$ | |
| **Occupation** |  |  |  |
| Working | 81.7 | 18.3 | 208 |
| Not working | 64.8 | 35.2 | 197 |
| | $\chi^2 = 19.9$ | $p=0.001$ | |
| **Wealth Index** |  |  |  |
| Poor | 64.3 | 35.7 | 132 |
| Middle | 76.0 | 24.0 | 108 |
| Rich | 79.2 | 20.8 | 165 |
| | $\chi^2 = 11.8$ | $p=0.016$ | |
| **Knowledge of any FP method** |  |  |  |
| Knows FP methods | 72.6 | 27.4 | 392 |
| Knows nothing | 98.3 | 1.7 | 13 |
| | $\chi^2 = 5.57$ | $p=0.000$ | |
| **Knowledge of fertile period (ovulation cycle)** |  |  |  |
| Don't know | 79.8 | 20.2 | 244 |
| Know (After periods) | 63.8 | 36.2 | 161 |
| | $\chi^2 = 17.1$ | $p=0.005$ | |

The prevalence of teenage pregnancy decreased with increasing education level, and the association was statistically significant (p=0.039). Teenage pregnancy in male headed households (31.7%) was statistically significantly different (p=0.000) from female headed households.16.2%). There was twice teenage pregnancy rate in male headed households compared to the female headed households. Teenage pregnancy in married (91.2%) and ever married (84.1%) teens was statistically significantly different (p=0.000) from those who were never married (8.9%).

Teenage pregnancy is statistically significantly associated with occupation of the teens (p=0.001). There was almost twice teenage pregnancy rate in the not working teens (35.2%) compared to the working class (18.3%). The results further indicate that teenage pregnancy decreased with increase in wealth index.

There were statistically significant associations between knowledge of family planning methods (p=0.000) and teenage pregnancy. In addition, there was a higher teenage pregnancy in teens who had knowledge of ovulation cycle (36.2%) compared to those who had no knowledge at all (20.2%), and the difference was statistically significant (p=0.005).

### Predictors of teenage pregnancy

The variables with significant associations with teenage pregnancy were included in the binary logistic regression model, and the results are presented in Table 3.

**Table 3:**
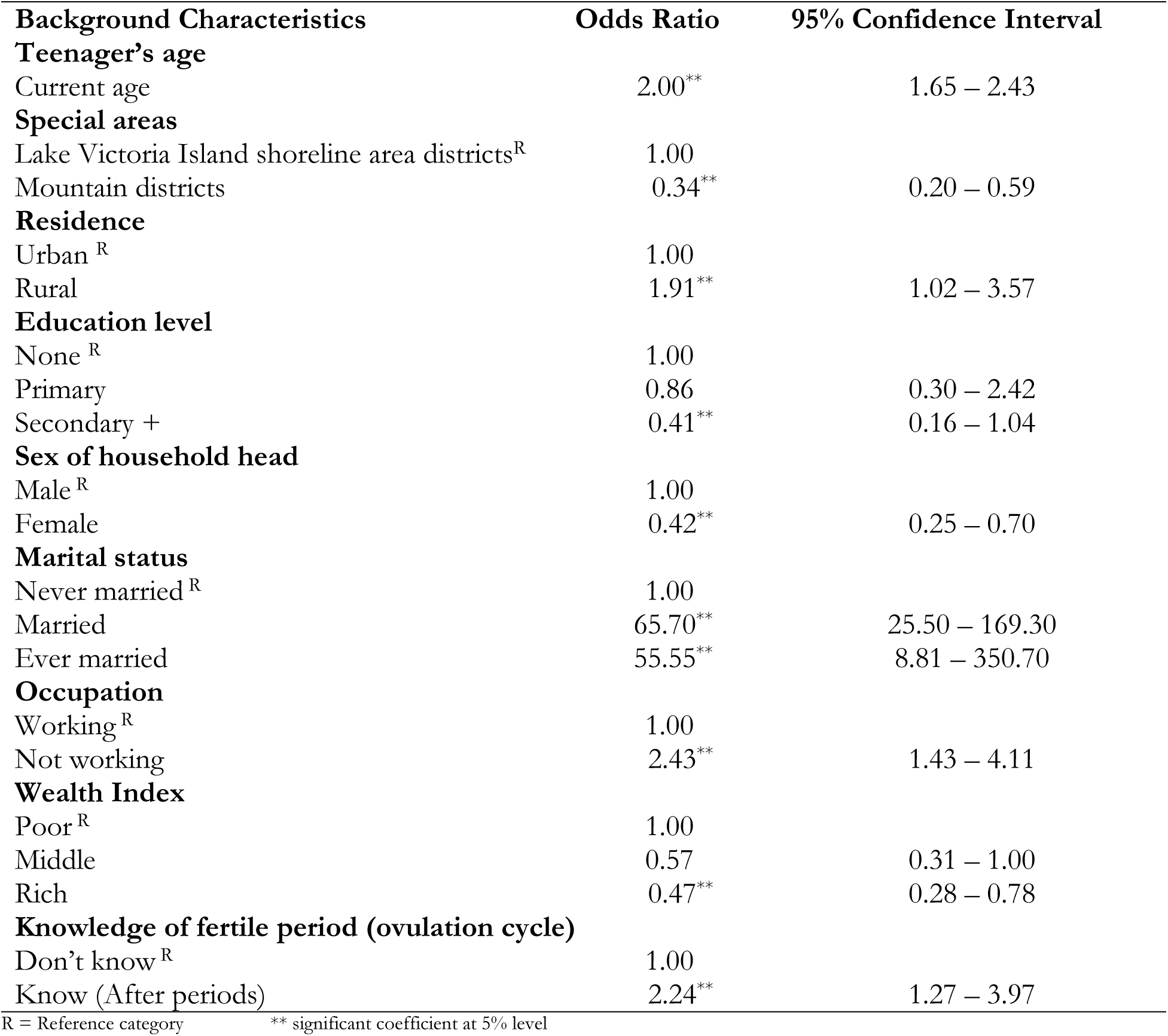
Predictors of teenage pregnancy

The logistic regression analysis results indicate that older teens were twice (OR=2.00, 95% CI =1.65 – 2.43) more likely to experience teenage pregnancy than the younger teens. This finding is consistent with the study conducted by (Kurth et al., 2010). Similarly, teens who lived in rural areas were almost twice (OR=1.91, 95% CI=1.02 – 3.57) more likely to experience teenage pregnancy than those who lived in urban areas. Also, teens from mountain districts had less odds. With regard to education level, compared to teens with no education, the risk of teenage pregnancy reduced among teens with primary education (OR=0.86, 95% CI=1.30 – 2.42), and reduced further among teens with secondary or higher education level (OR=0.41, 95% CI=0.16 – 0.70).

Table 3 further shows that sex of household head was a predictor of teenage pregnancy. Teens from households headed by a female were less likely (OR=0.42, 95% CI=0.25 – 0.70) to experience teenage pregnancy than their counterparts from male headed households. Marital status was a statistically significant predictor of teenage pregnancy. Married (OR = 65.70, CI= 25.50 – 169.30) or ever married (OR=55.55, CI=8.81 – 350.70) teens and very high odds compared to the never married teens. Furthermore, in comparison with working teens, those who are not working were more likely (OR=2.54, 95% CI= 1.43 – 4.11) to experience teenage pregnancy.

Results show that the socio-economic status of a teen was a predictor of teenage pregnancy. There were reduced odds of experiencing teenage pregnancy among teens with increasing wealth index. Teens who belonged to the rich quintile were less likely (OR=0.47, 95% CI=0.28 – 0.78) to experience teenage pregnancy than their counterparts belonging to the poor category. With regard to knowledge of the ovulation cycle, compared to teens who do not know, teens who know the ovulation cycle had over twice (OR=2.24, 95% CI=1.27 – 3.97) the risk of experiencing teenage pregnancy.

## Discussion

Teenage pregnancy in the Uganda Lake Victoria Island districts was 48.3% and 24.3% in the Mountain districts. Overall, teenage pregnancy in these special areas was 26%, which is slightly over the national figure of 24.8%. This finding is consistent with studies in other countries which show high teenage prevalence ranging from 20% to 50% (Manzi, Ogwang, Akankwatsa, Wokali, & Obba, 2018; Yakubu & Salisu, 2018) Omoro et al., 2017; Amoran, 2012; Lillian & Mumbango, 2015; Jewkes et al., 2001). This reflects a pattern of sexual activity which puts teenagers at a risk of HIV/AIDS. This could be attributed to poverty, peer pressure influence or lack of Information Education and Communication (IEC) materials to promote safe sex.

As observed by Omoro et al (2017), this study shows that teenagers who lived in rural areas were more likely to have teenage pregnancy compared to their counterparts in the urban settings. This is because teenagers from the rural areas are less educated and have limited access to sexual health services than their urban counterparts. This is an issue of concern given that UBOS (2016) reports that 85% of Uganda’s youth live in rural areas. Providers of rural health care services should make facilities youth friendly. Policies should be designed to promote youth involvement in safe sex activities.

Results indicate that teens with primary, secondary or higher education were less likely to have teenage pregnancy compared to those with no formal education. This could be due to lack of school fees and scholastic materials, as well as lack of transport to and from school as similarly observed by Omoro et al (2017). Education plays an important role in empowering teens with information and knowledge about safe sex, as reported in other studies(Cook, 2010; Gyan, 2013; Omoro et al., 2017; Stanger-Hall & Hall, 2011; Westway, Barratt, & Seeley, 2009).

Teens from female headed households were less likely to be exposed to teenage pregnancy compared to teens from male headed households. This could be due to lack of information communication about sexual reproductive health from the male heads of households. And as a result teens engage in early sexual activities that lead to teenage pregnancy. Girls tend to be much closer to female household heads than males household heads. Female household heads may easily discuss sexual reproductive health (SRH) issues with teenage girls than male household heads, a situation that would probably reduce the likelihood of early sexual relations and pregnancy. By contrast, the rapport between teenage girls and male household heads could be poorer and hence less discussion on SRH issues. However, Ellis et al (2003) observed that greater exposure to father absence was strongly associated with elevated risk for early sexual activity and adolescent pregnancy.

Wealth status was a significant predictor of teenage pregnancy. Teens from rich wealth quintile had reduced odds compared to those from the poor quintile. Teens from rich households are perceived to have access to desired basic childhood necessities like education and knowledge about sexual health care. However, in Nigeria, Amoran (2012) observed that students are exposed to teenage pregnancy because of low socio-economic status of the households.

## Conclusions

Increased age, rural residence, secondary or higher education level, occupation, and socio-economic status were found to be statistically significant predisposing factors of teenage pregnancy in Uganda Lake Victoria Island shoreline area and mountain districts. We recommend that teenagers should be provided with sexual education for them to learn about the changes they go through and their sexual reproductive health rights. Other measures such as promoting household wealth creation and ensuring girls keep in school by providing them with scholastic materials and other school requirements. Also, provision of teenage-friendly health services at health facilities that include a wide range of options, as well as medically accurate counselling and information could mitigate teenage pregnancy.

